# Improved prediction of site-rates from structure with averaging across homologs

**DOI:** 10.1101/2024.02.27.582061

**Authors:** Christoffer Norn, Fábio Oliveira, Ingemar André

## Abstract

Variation in mutation rates at sites in proteins can largely be understood by the constraint that proteins must fold into stable structures. Models that calculate site-specific rates based on protein structure and a thermodynamic stability model have shown a significant but modest ability to predict empirical site-specific rates calculated from sequence. Models that use detailed atomistic models of protein energetics do not outperform simpler approaches using packing density. We demonstrate that a fundamental reason for this is that empirical site-specific rates are the result of the average effect of many different microenvironments in a phylogeny. By analyzing the results of evolutionary dynamics simulations, we show how averaging site-specific rates across many extant protein structures can lead to correct recovery of site-rate prediction. This result is also demonstrated in natural protein sequences and experimental structures. Using predicted structures, we demonstrate that atomistic models can improve upon contact density metrics in predicting site-specific rates from a structure. The results give fundamental insights into the factors governing the distribution of site-specific rates in protein families.

## Introduction

There is a substantial difference in the overall rate by which different proteins evolve, but there is also a great variation in the rate of evolution at different positions or sites within a protein. Site-specific rate variation carries information about the various fitness constraints acting on different parts of a protein, which makes them interesting probes of evolution. These constraints can be divided into functional constraints, which relate to features that directly affect the function of a protein, such as active sites of enzymes, and structural constraints^1^. For most positions, the local structure imposes the dominant fitness constraint since many proteins must maintain their fold to carry out their function and because misfolded species can be toxic^1-3^. By developing biophysical models that predict rate variation at sites in proteins from the structure, it is possible to separate rate variation due to functional and structural constraints^4^. In general, it is observed that sites that are buried on the core of proteins are more conserved than surface-exposed positions^5-8^. Biophysical models directly and indirectly built on protein stability fitness constraints have been developed to understand rate variation in proteins. They can directly model the change in stability based on predicted changes in free energy (ΔΔG) upon mutation^3, 4, 9^. Alternatively, they can indirectly account for stability by using solvent accessibilit^5-8^ (that relates to the cost of burial of hydrophobic residues, like Residual Surface Area (RSA)) and packing density at sites^4, 9-13^ (that relates to the mean interaction energy of a site with other residues in a protein, like Weighed Contact Number (WCN) ^11^). The indirect and direct models have been shown to perform on par in terms of their ability to explain site-specific rate variation^4^, although a substantial level of detail is lost when features such as solvent accessibility and packing density are used to model the heterogeneity in rates within a protein. Each site evolves within a specific chemical and structural environment that can vary substantially for sites with similar solvent exposure or packing density.

The relative success of the more coarse-grained models of protein energetics can partially be understood by considering how empirical site-rates are calculated. Site-specific rates are inferred using multiple sequence alignments and a phylogenetic tree. A single substitution rate-variable is assigned to each site in the protein and inferred with Bayesian^14^ or maximum likelihood approaches^15^. Hence, the site-specific rates reflect an average over the different branches of the tree. A protein structure corresponds to a single leaf in the phylogenetic tree, representing a specific structural environment sampled in evolution. Coarse-grained metrics like solvent exposure and packing density may better represent an average state of a site during evolution than a single structure.

Here, we hypothesize that it should be possible to account for the effect of tree-averaging by calculating rates based on several structures of leaf sequences in a phylogenetic and averaging the results. We show that site-specific rates from different homologous structures vary substantially and that averaging over site-specific rates substantially improves the ability to predict rate variation. When using structures from simulated evolutionary trajectories - where the fitness function, evolutionary dynamics, and phylogeny is known - the prediction of site-specific rates is near perfect after averaging. The same effect is seen with experimental structures, but the resulting correlations do not improve beyond contact density as predictors of site-specific rates. This, however, can be achieved by using predicted protein structures rather than experimental structures as the basis for rate estimations.

## Results

### Prediction of site-specific evolutionary rates based on crystal structures

Although the three-dimensional structure of proteins evolves slowly, the microenvironment in sites can vary substantially between homologous proteins. This could result in significant variation in evolutionary rates among corresponding sites within a protein family. To study this effect we calculate site-specific evolutionary rates using homologous crystal structures from the same protein family. Site-specific rates are calculated using the TMS (Thermodynamic Mutation-Selection) model. TMS is based on the mutation-selection model^3^ and calculates site-specific rates using a stability fitness model where thermal stability values are estimated from structure-based ΔΔG calculations from Rosetta. We generated multiple sequence alignments for 6 protein families for which multiple (11-19) experimental structures of homologs have been determined. For each structure, we calculate the site-specific rates and correlate them with empirical rates estimated for that protein family estimated from sequence and phylogeny using LEISR^15^, **Fig. 1**.

**Fig. 1.**
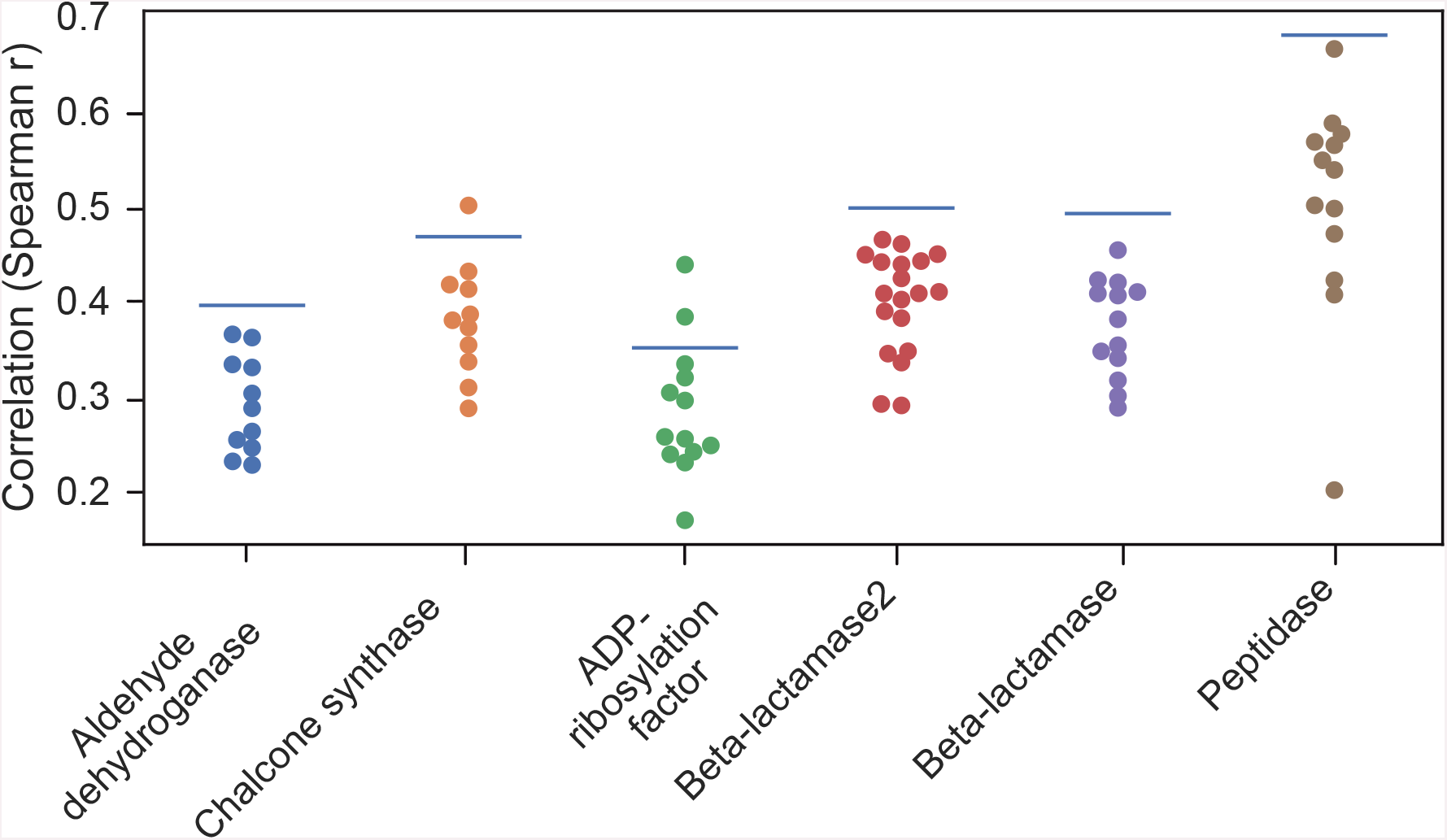
Distribution of correlations between site-specific rates predicted from structures and empirical rates. Results for 6 different protein families where each point represents a crystal structure used to calculate site-specific rates. Averaged site-specific rates indicated as blue bars.

The results show a substantial variation between protein families with correlation values ranging from 0.3 to 0.5. But there is also significant variation within each family. For the peptidase family, the correlations range from 0.20 for one structure to 0.67. The substantial variation in correlation with empirical rates for structure-based models is consistent with previous results, showing that between 10 to 60% of rate variation can be explained from the structure using simple metrics like solvent exposure and packing density^9^. The results indicate that the structural environment in each protein can be substantially different, resulting in rate variation across homologous. However, one caveat is that ΔΔG predictions have limited accuracy and the variation in correlation with empirical rates could reflect misestimated stability values.

### Experimental ΔΔG measurements only result in minor improvements to rate predictions

To assess the impact of errors in predicted ΔΔGs, we predicted the site-specific rates for the small domain from protein G, GB1, for which nearly all point mutations were recently characterized using chemical denaturation^16^, excluding only cysteines and tryptophan at one site used for detection. There is a small improvement in using rates estimated from experimental ΔG values compared to values predicted from ΔΔG calculations with Rosetta, with correlation values increasing from 0.56 to 0.58. Nonetheless, this result suggests that highly accurate ΔΔG values may not be sufficient criteria to get high correlations with empirical site-specific rates. Regardless, the computational prediction of ΔΔG values will reduce correlation with empirical site-specific rates. ΔΔG calculations can be sensitive to the resolution and quality of crystal structures, even after preparative energy refinement. We included several independently solved crystal structures of the same proteins in Fig. 1. For example, there are three structures of the protein ficin (PDB ID: 4YYQ, 4YYR, 4YYW). The correlations vary from 0.55 to 0.67 for these structures, indicating that relatively small changes in 3D coordinates can have a substantial impact on the calculated rates. A caveat with the correlation analysis is the limited number of homologous sequences (32) available for empirical site-specific rate calculations of GB1, making the empirical rate values more uncertain.

### Simulation of protein evolution using an all-atom simulator

To better understand the factors causing variations in correlations with with site-specific rates calculated from homologous structures, it is helpful to use simulated data where we can avoid uncertainty due to uncertainties in the true 3D coordinates, ΔΔG–values, phylogeny, variation in fitness model over time and empirical site-specific rates. We have recently developed a method called RosettaEvolve^17^ that enables us to simulate the evolution of a protein along a phylogeny using a stability-based fitness model and an atomistic energy function. Starting from a defined phylogenetic tree, evolutionary trajectories are simulated along branches. Random mutations are introduced at the DNA level, using the same model employed in TMS, with substitution probabilities controlled by parameters κ (transition-transversion bias) and ρ (whole codon mutation rate), and with fixation probabilities calculated using ΔG values from predicted ΔΔG values and assumed ΔG_nat_ together with a value for the effective population size N_e_. Limited changes in backbone structure occur during the simulation as backbone adapts to introduced mutations.

### Averaging over predicted rates from simulated trajectories results in near perfect recapitulation of site-specific rates

We estimated phylogenetic trees from natural sequences for four small protein domains and used those together with experimental structures to simulate multiple sequence alignments. We can track the evolutionary process in the simulation so that the true average site-specific rate for each site in the protein can be measured. The instantaneous rate is not constant across evolutionary time but fluctuates with the current stability of the protein^17^. The rate measured by counting the number of substitutions at each site in the protein across all branches is an average across evolutionary time. Since the same energy function used to simulate the sequences is also used to estimate site-specific rates, there is no uncertainty in the energy calculations, and there is no uncertainty over the true structure of the sequences. Furthermore, the fitness function is constant across all branches and the assumptions in the TMS model are compatible with those in the evolutionary dynamics simulation. In Fig. 2 we have calculated correlations between site-specific rates from the TMS model and empirical site-specific rates calculated from the simulation of many structures at the leaves of the tree.

**Fig. 2:**
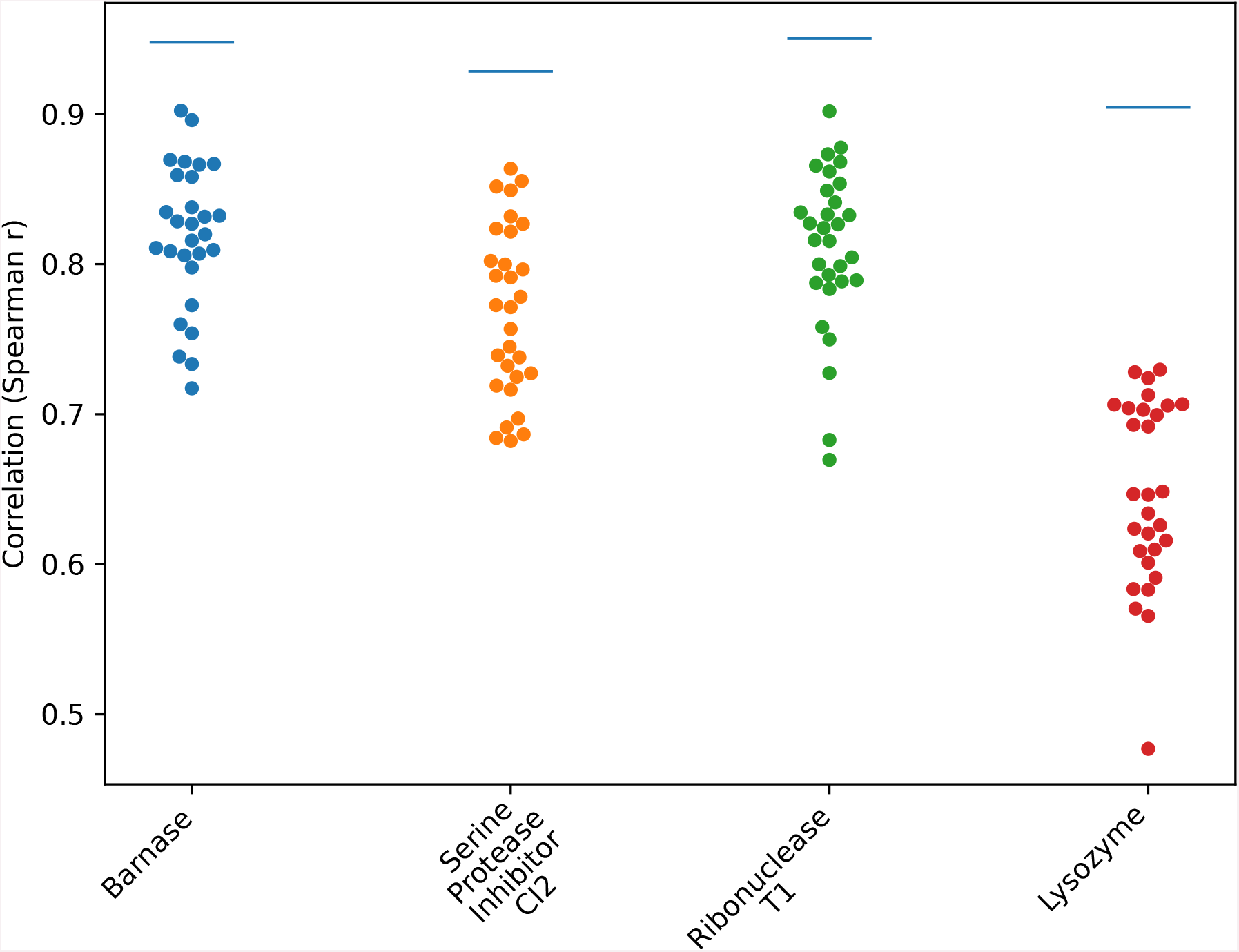
Correlations between site-specific rates estimated from sequences simulated by RosettaEvolve and empirical site-specific rates. Simulations based on four small single-domain proteins with PDB codes 1a2p (Barnase), 2ci2 (Serine Protease Inhibitor CI2), 1rn1 (Ribonuclease T1), and 1el1 (Lysozyme). Spearman r correlation values with empirical site-specific rates calculated from the RosettaEvolve trajectories. Averaged site-specific rates indicated as blue lines.

There is significant variation in the correlations within each family of evolved structures. A packing density metric, Weighted Contact Number (WCN)^13^ has a substantially lower correlation with the empirical rates, with correlations between 0.46-0.51, compared to energy-based rates. Due to the limited variation in backbone coordinates during the simulation, the WCN is very similar across the structural ensemble of each family.

The variation in correlation with empirical site-rates in Fig. 2 is the result of calculating rates from protein structures with very similar backbones, but different sequences. Hence, site-rate variations in these systems are caused by differences in the energetic environments and overall stability of the homologs. The empirical rates represent an average across the phylogenetic tree, estimated from sequences sampled at the leaves of the tree (or counted substitutions). This suggests that an averaging of the predicted site-specific rates should improve the correlation with the empirical rates. In Fig. 2 we show the effect of averaging site-specific rates from simulated alignments. For all four systems site-specific rate averaging recovers the empirical rate accurately, with Spearman r values of 0.90-0.95. In all cases the averaged correlation is an improvement on all the individual predictions. Averaging the site-rates before correlation with sequence-based rates is substantially better than averaging the correlations of the individual protein structures.

In Fig. 1 we apply the same averaging on the site-specific rates estimated from crystal structures within the same protein family. In 4 out of the 6 cases, the weighted averaged values improve over all the individual correlation calculated with the individual structures from the same family. In all cases averaging the site-specific rates improves upon the average correlation calculated from individual structures. This demonstrates that averaging site-specific rates across multiple structures leads to substantial improvements over using a single experimental structure.

### Improved correlations with models

We can now compare correlations calculated with a packing density metric, WCN, which has been shown to have the best performance for site-rate prediction. The average correlation with WCN across the 6 families is 0.52 while for ΔΔG-based rate prediction it is 0.48. So, although averaging the site-specific rates across the homologous proteins improves the correlations, it does not improve over WCN. One remaining source of error is the uncertainty of the true structure of experimental structures, where small differences in the atomic coordinates can give rise to substantial differences in predicted ΔΔG values. To investigate this factor, we predicted the structure of sequences using the deep learning structure prediction method trRosetta^18^ and used those models as the basis for ΔΔG predictions. While trRosetta models may not have the correct local structure at every site in the protein, it produces a consistent way of generating the 3D coordinates across homologous sequences. While the crystal structures were energy refined with Rosetta before ΔΔG predictions, trRosetta models are likely to produce structures that are more consistent with the Rosetta energy function used for rate calculations. We modeled the 7 previously described protein families using this approach. As with the other examples, site-rate averaging improves the correlation to empirical rates considerably, Fig. 3.

**Fig. 3:**
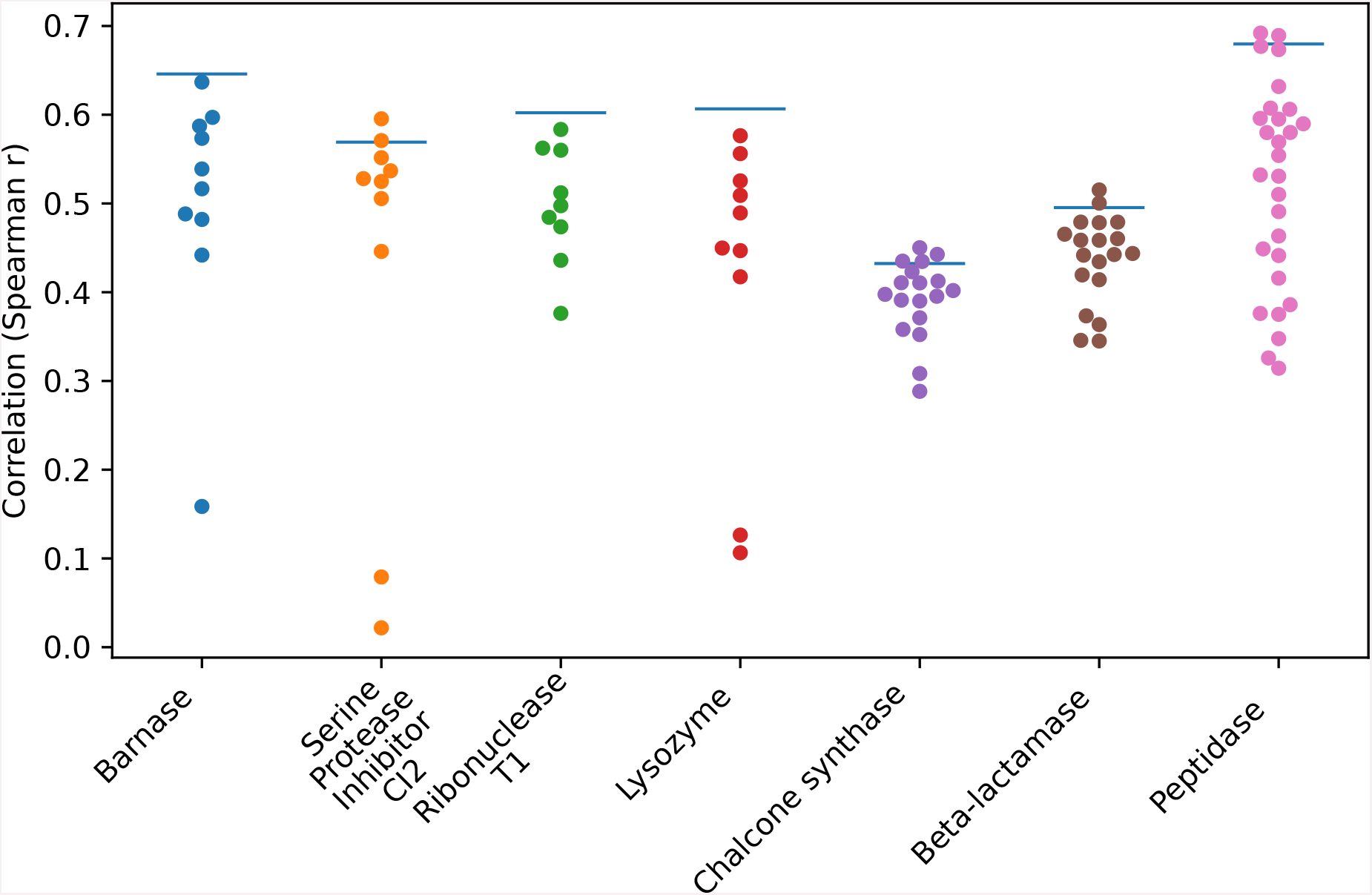
Correlations between site-specific rates estimated from predicted structures and empirical site-rates. Site-specific rates were estimated from sequences homologous to the systems used in the RosettaEvolve benchmark (Barnase, Serine Protease Inhibitor CI2, Ribonuclease T1, and Lysozyme) and sequences used in the crystal structure benchmark. Spearman r correlation values with empirical site-specific rates calculated from the RosettaEvolve trajectories. Averaged site-specific rates indicated as blue bars.

In Fig. 4, the correlations with empirical site-specific rates are compared with the results using WCN to predict site-specific rates. There is now an improvement over WCN of using the energy-based site-specific rate calculation, even though the correlations are not as high as observed in the simulated data.

**Fig. 4:**
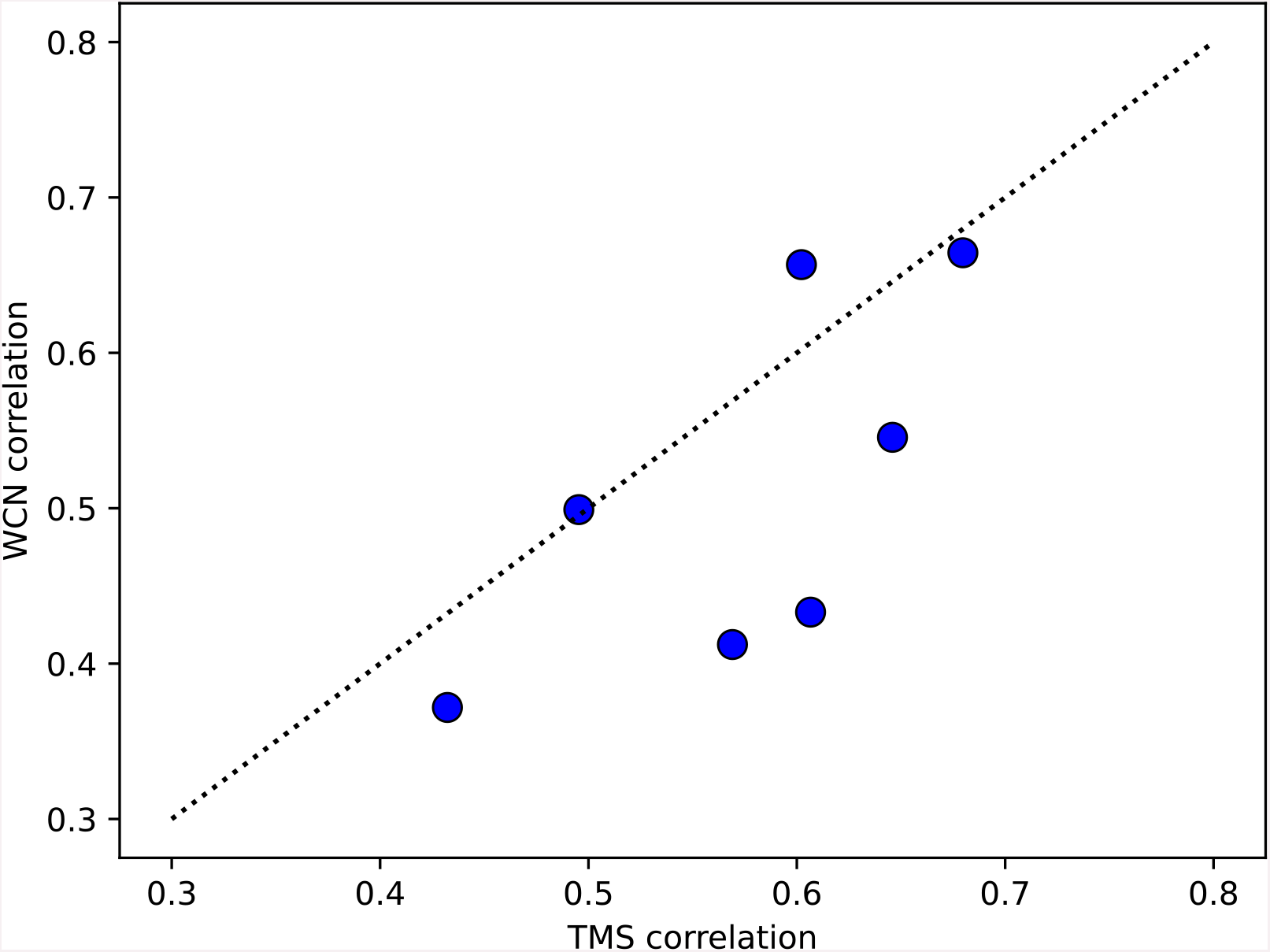
Comparison of site-specific rates predicted with the TMS model and WCN. Site-specific rates averaged across multiple models predicted with trRosetta for each of the 7 families in Fig. 3.

## Discussion

The fold of a protein generates a three-dimensional context for sites that is similar across homologs. Nonetheless, amino acid substitutions occur at sites that can have substantially different contexts in terms of atomic interactions within the background of each protein sequence. This results in epistasis^19^, where the impact of substitution can be very different in one sequence environment compared to another even with almost identical overall structures. We can thus expect that a model that considers the detailed atomic and energetic context should be able to predict site-specific rates with higher accuracy. However, models based on packing density (WCN), that blur out atomic details, have equal performance to atomistic methods. Echave et al. developed a method for site-specific rate prediction based on ΔΔG predictions with Rosetta and FoldX or a *v*stress-based*v* energy model using WCN and demonstrated similar performance using either approach^9^.

A fundamental issue in using a single structure and a ΔΔG-based model to predict site-specific rates is that empirical site-specific rates report an average across all branches across the phylogenetic tree. The homologous proteins have sampled a range of structural and energetic environments resulting in substitution patterns observed in extant proteins. Simulations have demonstrated that how well accommodated an amino acid at a site is not constant over time, because the environment adapts to the presence of new amino acids^20, 21^. Evolutionary dynamics simulations, using energy models based on covariation analysis^22^ as well as atomistic energy functions^17^, have also demonstrated how site-specific rates fluctuate over time or phylogenetic branches (heterotachy). Fluctuations in site-specific rates can be largely coupled to variations in overall protein stability^17^, resulting in site-specific rates fluctuations that change around an average rate[17]. Taking this averaging into account is critical when comparing it to empirical values. We have previously used the TMS model to predict amino acid substitution rates with averaging across sites^17^. We observe that the TMS explains around 65% of the variation of amino acid substitution rates averaged across sites (as captured in substitution matrices LG^23^ and WAG^24^). This performance is, to a large extent, the consequence of the removal of variations of the details of individual sites in the site-averaged data but highlights the ability of the stability fitness model in explaining average substitution behavior in proteins. The fact that empirical site-specific rates are the result of averaging across many microenvironments explains why WCN is a comparably good predictor of site-specific rates even though it removes many of the atomic details. Nonetheless, we should still expect an energy-based model to have a higher ceiling in terms of the ability to predict site-specific rates.

Our RosettaEvolve simulations, where uncertainty in atomic coordinates, stability calculations, phylogeny, fitness function and empirical site-specific rates can be disregarded, show that the TMS model averaged across proteins can fully recover the true rates to a substantially higher degree than WCN.

For natural sequences with known structures, we also demonstrated the benefit of averaging across predicted site-specific rates. Nonetheless, the correlations are substantially lower than for simulated evolution. This can be a consequence of more complex fitness pressures beyond stability acting at sites in proteins. But it can also be due to uncertainties on protein stability estimations and noise in experimentally determined protein structures. The improvement of results with predicted models of protein structure over experimental structures suggests that it is more important to provide a consistent method for site-rate averaging than to have the exact coordinates, as it properly averages out structural noise.

The results using predicted structures point towards using them in the future. Structure predictions based on deep learning methods have become very accurate in predicting the structure of proteins ^25-27^. Thus, we can generate structures for extant sequences in a protein family and use them for site-specific rates prediction.

A primary application of site-specific rates based on a stability fitness model is to use them as baselines for identifying functional sites evolving with additional fitness pressures beyond stability. But the results here also provide fundamental insights into the factors shaping the distribution of empirical site-specific rates.

## Methods

### Site-specific rate prediction from protein structure

A detailed description of the TMS model can be found in Norn et al. ^3^. Briefly, the model is based on the mutation-selection framework described at the codon level. The relative instantaneous rate between (*q*_*uv*_) from codon *u* with amino acid *i* to codon *v* with amino acid *j* is calculated as the product of the proposal rate (*p*_*uv*_) for mutating codon *u* to *v* and the fixation probability (*f*_*uv*_), *q*_*uv*_ = *p*_*uv*_ · *f*_*uv*_. The proposal rate is evaluated using a nucleotide substitution model K80 with a parameter (κ) describing the relative rate of transitions to transversions, and whole codon mutation rate (ρ). The fixation probability for codon substitutions was estimated using Kimura’s fixation probability equation^28^ for a diploid organism (parametrized by the effective population size N_eff_) and a stability fitness model^29^. Thermodynamic stability of amino acid variants was calculated using assumed stability of the wildtype sequence (ΔG_nat_) and ΔΔG predictions using Rosetta, as described in Norn et al.^3^. Site rates were calculated from the instantaneous rate matrix constructed from the rates of all codon substitutions at a given site by summing the flux between all connected codons *u* and *v* that encode the same amino acids. The parameters for the TMS model (κ=2.7, ρ=0.1, and N_e_=10^2.2^) were selected by optimizing the correlation between empirical site-rates calculated by Rate4site^14^ and the site-rates calculated from TMS on the benchmark set of 66 described in Norn et al.^3^. The optimal ΔG_nat_ will vary between each structure and can in principle be optimized for the highest correlation with sequence-based site-rates. In this study the value was fixed and set to corresponding to 6.25 kcal/mol. This value is close to the average value when the correlation for each protein structure is optimized independently. For the other (smaller) systems studied in this work, including structures from RosettaEvolve trajectories, a value of 5.4 kcal/mol was used. Sites with cysteines were excluded due to challenges with dealing with disulfide formation in the energy calculations. Site rates were also compared with WCN using the CA packing density using a script presented by Jack et. al.^30^.

### Preparation of structures

The crystal structures used in this work were energy refined using the Rosetta macromolecular modeling package^31^ using the method described by Nivon et al^32^. before use in stability calculations or evolutionary simulations. Prediction of ΔΔG values was carried out as described in Norn et al.^3^.

### Empirical rate calculations and correlations with calculated rates

Site-rates were calculated from sequences and phylogenetic trees using LEISR^15^. Natural sequences for empirical site-rate calculations for the systems studied by RosettaEvolve were harvested using Consurf^33^ followed by the generation of phylogenetic trees using IQ-TREE^34^. The sequences and multiple sequence alignments for the crystal structure benchmark were generated using the uniclust sequence database^35^ and hhblits^36^ and based on pfam families^37^ (ADP Ribosylation factor: Arf, Beta-lactamase-2: Beta-lactamase2, Beta-lactamase: Beta-lactamase, Peptidase: Peptidase_C1, Chalcone synthase: Chal_sti_synt_N_66, Aldehyde dehydrogenase: Aldedh). Initial estimates of site-specific rates for these protein families were done with Rate4site^14^, but the empirical rates correlated to calculated rates were estimated from LEISR using the tree from Rate4site.

When correlating empirical rates from LEISR with the calculated rates from TMS, sites with high uncertainty in estimated site-specific rates were excluded (upper and lower ML rate estimate bigger than 5 units), and sites with outliers (ML site-specific rate estimate < 0.0001 and ML site-specific rate estimate > 5).

### Rates from experimental experimental stability values

Experimental stability values for protein GB1 were taken from values deposited by Nisthal et al.^16^ into the ProtaBank database^38^. There are 1120 single mutant data for protein GB1, and for 157 mutations data is missing. Missing data was given a stability of 1000 kcal/mol in the calculation. The sequences for sequence-based site-specific rates were collected using a previous version of the Consurf^33^ server with two sequences from the protein data bank manually added (ID 1QKZ and 1IGD). The phylogenetic tree was generated with IQ-TREE^34^ and LEISR^15^ was used for site rate calculation. The rate calculation with TMS was based on the crystal structure with PDB ID 1PGA^39^.

### RosettaEvolve trajectories

Simulated sequence alignments were generated with RosettaEvolve using the approach described in Norn et al. for the proteins with pdb id (1a2p, 1rn1, 2ci2, and 1el1). Simulations were done with κ=2.7 and ρ=0.1, N_eff_=10^4.4^. Several different evolutionary trajectories were simulated for each protein with different assumed stabilities of the protein, controlled by the offset parameter described in Norn et. al.^17^ that is combined with the Rosetta energy of the wildtype sequence to define the stability of the protein. The data in this study used the following offset parameters: 1a2p: -314.299, 1rn1: -210.276, 2ci2:-143.898, 1el1: -141.062. Phylogenetic trees used for the simulations were calculated using RAxML^40^ based on sequence alignments generated by Gremlin^41^. Before simulating phylogenetic trees using RosettaEvolve, the structures were equilibrated using RosettaEvolve in a simulation along a single branch as described in Norn et al.^17^.

### Rate averaging

Averaged rates were calculated with a simple averaging of site-rates from the instantaneous rates estimated from each structure in the set. A minimal number of 5 structures are required to be included for a site to be included in the analysis (excluding positions in the MSA with few sequences) and the protein was required to have at least 50 contributing sites for the rate correlation.

For the RosettaEvolve sequence only a subset of structure at the leaves was used for rate averaging. To get a representative sample two approaches were used. In the first the phylogenetic tree used to simulate sequences was analyzed with BranchManager^42^ and sequences were sampled according to leaf weighs. In the second approach, we used a method that prunes the tree of the leaf nodes with the shortest branch lengths until you get a tree with the number of requested leaf sequences. The final selection sequence/structures for averaging was as a combination of both.

## Data availability

Data for this study can be found at https://github.com/Andre-lab/Site-rate-averaging.

